# Aging reduces motivation through decreased *Bdnf* expression in the ventral tegmental area

**DOI:** 10.1101/2023.01.19.524624

**Authors:** Hanyue Cecilia Lei, Kyle E. Parker, Chao-Cheng Kuo, Carla M. Yuede, Jordan G. McCall, Shin-ichiro Imai

## Abstract

Age-associated reduced motivation is a hallmark of neuropsychiatric disorders in the elderly. In our rapidly aging societies, it is critical to keep motivation levels high enough to promote healthspan and lifespan. However, how motivation is reduced during aging remains unknown. Here, we used multiple mouse models to evaluate motivation and related affective states in young and old mice. We also compared the effect of social isolation, a common stressor in aged populations, to those of aging. We found that both social isolation and aging decreased motivation in mice, but that *Bdnf* expression in the ventral tegmental area (VTA) was selectively decreased during aging. Furthermore, VTA-specific *Bdnf* knockdown in young mice recapitulated reduced motivation observed in old mice. These results demonstrate that maintaining *Bdnf* expression in the VTA could promote motivation to engage in effortful activities and potentially prevent age-associated neuropsychiatric disorders.

## Introduction

In 2017, the World Health Organization reported that more than 20% of people aged 60 and over have a psychiatric or neurological disorder, including dementia, depression, and anxiety^1^. Older adults are also more susceptible to stressors such as social isolation, a common risk factor for many mental disorders and health issues particularly during the COVID-19 pandemic^2–4^. These age-associated psychiatric conditions are quite often associated with reduced motivation to engage in effortful activities^5^. Furthermore, maintaining high motivation levels is indicative of healthy mental conditions in old age^6^. How aging impacts mental functions, particularly motivation, and the underlying biological mechanisms, however, remains largely unknown.

To determine the neurobiological mechanisms of aging-induced decreased motivation, we adapted multiple mouse models thought to impact motivation. In particular, we used social isolation as a stressor to mimic a real-life condition in aging human population^7, 8^. We used sexually matured young adult (4 month-old) and old but still healthy (18 month-old) male mice. After one month of single or grouped housing, these mice were tested in a battery of motivational assays.

## Results

### Chronic social isolation and aging differentially impair motivational behaviors

Voluntary wheel-running activity (VWR) is innately rewarding to rodents even with no other extrinsic reward^9, 10^. Therefore, to test whether age impacts this endogenously motivated behavior, we compared VWR activities between young and old mice under grouped or socially isolated housing conditions. Old mice ran much less than young mice during the dark time, as reported previously^11, 12^. However, social isolation substantially reduced dark-time wheel-running in both young and old mice (Fig. 1a and b). Most VWR reduction occurred in the early dark time when mice reached their daily peaks of running (Extended Data Fig. 1a and b), and the effect of social isolation was greater in old mice, compared to young mice (40.3% vs. 25.7% reduction). Although VWR is a complex behavior that can be affected by both physiological (e.g., reduced skeletal muscle strength and energy metabolism) and motivational changes, these results suggest that both aging and social isolation undermine motivation.

**Fig. 1.**
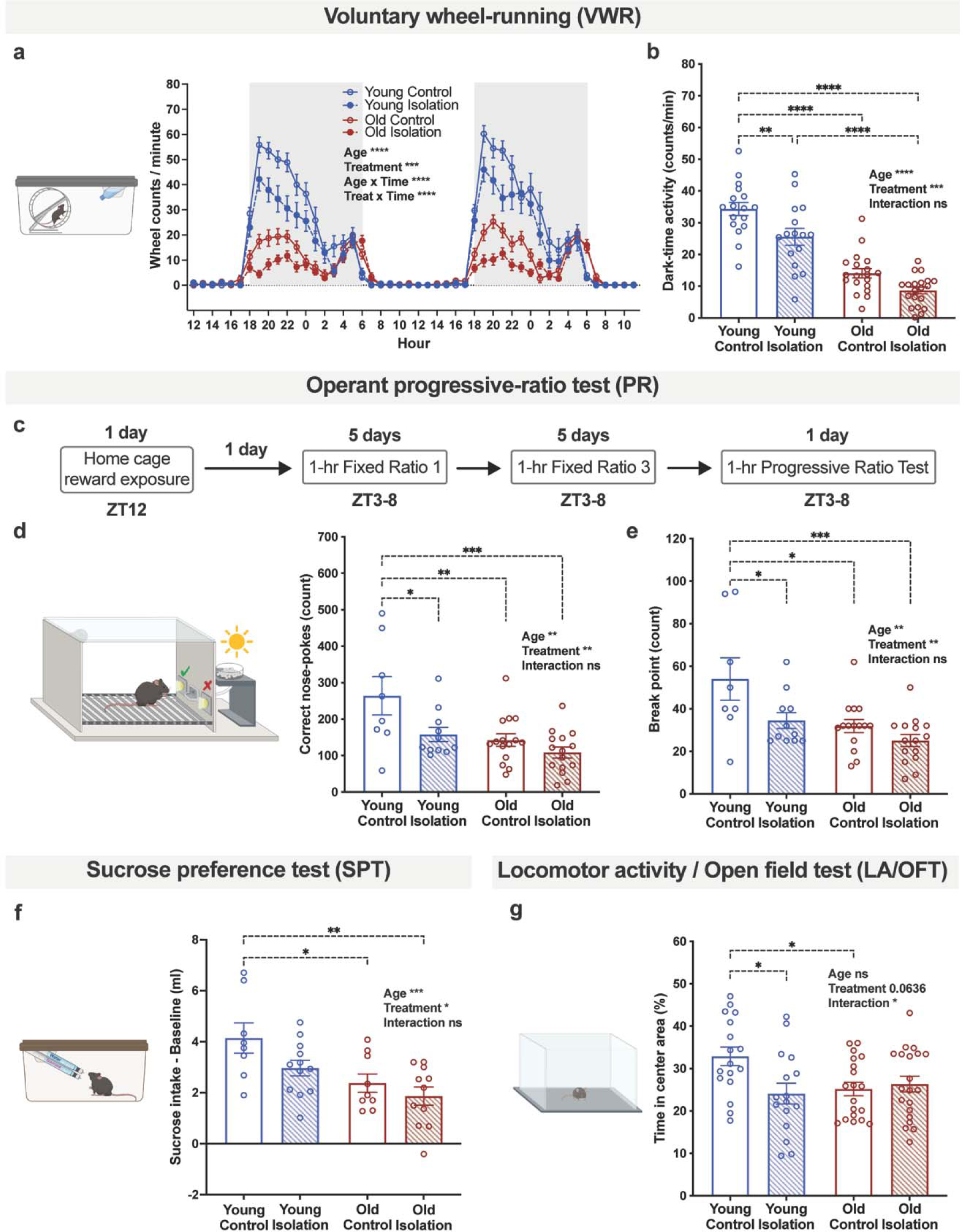
Chronic social isolation and aging differentially impair motivational behaviors and induce depressive- and anxiety-like behaviors. Young (5-7 months) and old (19-21 months) mice were assessed by batteries of behavior tests after a month of social isolation or grouped housing. (**a**) Voluntary wheel-running activity recording in consecutive 48 hours, shaded area representing the dark phase. (**b**) Average VWR activity during the dark phase (ZT12-0). (**c**) Calendar illustrating the operant progressive ratio test protocol. Correct nosepokes performed (**d**) and break points reached (**e**) in the PR test. (**f)** Delta of sucrose solution intake to baseline water intake in the sucrose preference test. (**g**) Percentage of time spent in the center area in the 1-hour locomotor activity and open field test (LA/OFT). Data were analyzed by 3-way ANOVA (**a**) and 2-way ANOVA (**b**, **d**, **e**, **f**, **g**) with *post-hoc* multiple comparisons. n=7-20. Error bars indicate SEM. Significant pair-wise comparisons and main effects of factors and interactions are labeled on the graphs: *p≤0.05; **p≤0.01; ***p≤0.001; ****p≤0.0001.

To directly examine whether aging and social isolation reduce motivation levels in mice, we next conducted an operant progressive ratio (PR) test to measure the amount of effort that animals are willing to expend to gain a palatable food reward. We habituated mice to sugar pellets in home cages two days before training and trained under fixed ratio-1 and fixed ratio-3 for five days each (Extended Data Fig. 1c). Following this task learning, we next tested the mice under a progressive ratio (PR) for one day (Fig. 1c). In the PR test, socially isolated young mice performed significantly fewer correct nose-pokes than group-housed controls (Fig. 1d). Interestingly, group-housed old mice showed significantly lower correct nose-pokes than group-housed young mice, but very similar to those of the socially isolated young mice (Fig. 1d). Socially isolated old mice tend to show further reduction in correct nose-pokes (Fig. 1d). Breakpoint analysis, which determines the maximum number of nose-pokes that an animal achieves to gain a single reward^13^, further demonstrates consistent reduction in effort to achieve rewards in old and socially isolated mice (Fig. 1e). In both socially isolated groups, there were tendencies of fewer entries into the food tray (Extended Data Fig. 1d). These results clearly suggest that reduced motivation is a shared outcome of social isolation and aging, and old mice seemingly responded less to the stress of social isolation perhaps due to the already reduced effort induced by aging.

### Both social isolation and aging induce depressive- and anxiety-like behaviors

Apathy and anhedonia are often comorbid with decreased motivation^14^. To determine whether normal reward processing is impacted by social isolation and aging, we next conducted the sucrose preference test (SPT). All groups of mice manifested robust preference to sucrose solution over water in the sucrose preference test (Extended Data Fig. 1e), indicating no obvious anhedonia. Nonetheless, the amount of sucrose solution consumed was selectively reduced by social isolation and aging (Fig. 1f), although the extent of reduction by social isolation did not reach statistical significance in either young or old mice. Reduction in energy demand was an unlikely explanation because daily food intake was not different for any group on the test day (Extended Data Fig. 1f). Notably, the results from the SPT showed very similar patterns to those from the PR test, supporting the notion that social isolation and aging both reduce motivated behavior in mice.

Finally, to determine whether approach-avoidance conflicts that underly some features of an anxiety-like state might be altered by social isolation and aging in young and old mice, we tested mice in the locomotor activity/open field test (LA/OFT) and elevated plus maze (EPM). Socially isolated young mice spent less time exploring the center area than the group-housed controls, whereas both group-housed and socially isolated old mice exhibited similarly lowered center time and distance traveled (Fig. 1g and Extended Data Fig. 2a and b). Contrarily, socially isolated young mice showed significant increases in time and distance traveled to explore the peripheral area (Extended Data Fig. 2a and b). Additionally, in the EPM, social isolation increased distance traveled and percentages of moving times in the close arm in both young and old mice, as we observed previously^15^, whereas there were no differences between young and old mice (Extended Data Fig. 2c-e). We also conducted the hole-board olfactory preference test and the social interaction test to assess other types of motivation-related social behavior between young and old mice in both group-housed and socially isolated conditions (Extended Data Fig. 3). Social isolation had no effect on either assay, except for a trend for isolated young mice to seek more social cues. Therefore, when motivation was reduced by social isolation and aging, certain anxiety- or depression-related behavioral states were also present. Together, these results suggest that both aging and social isolation reduce motivation in mice, but perhaps not additively.

### Aging impairs dark time-dependent enhancement of reward-seeking motivation

To determine the extent to which age alters motivation, we tested a new cohort of young and old mice across different testing conditions. Remarkably, we found that when food-deprived and tested during the dark time (Fig. 2a), differences in correct nose-pokes detected in the PR test between young and old mice were greatly enhanced (Fig. 2b). Similar results were obtained for break points (Fig. 2c). These results suggest that the reward-seeking motivation of young mice is enhanced by fasting and heightened during the dark time, whereas old mice fail to show such dark time-dependent enhancement of reward-seeking motivation, confirming that aging is a critical contributor to reduced motivation.

**Fig. 2.**
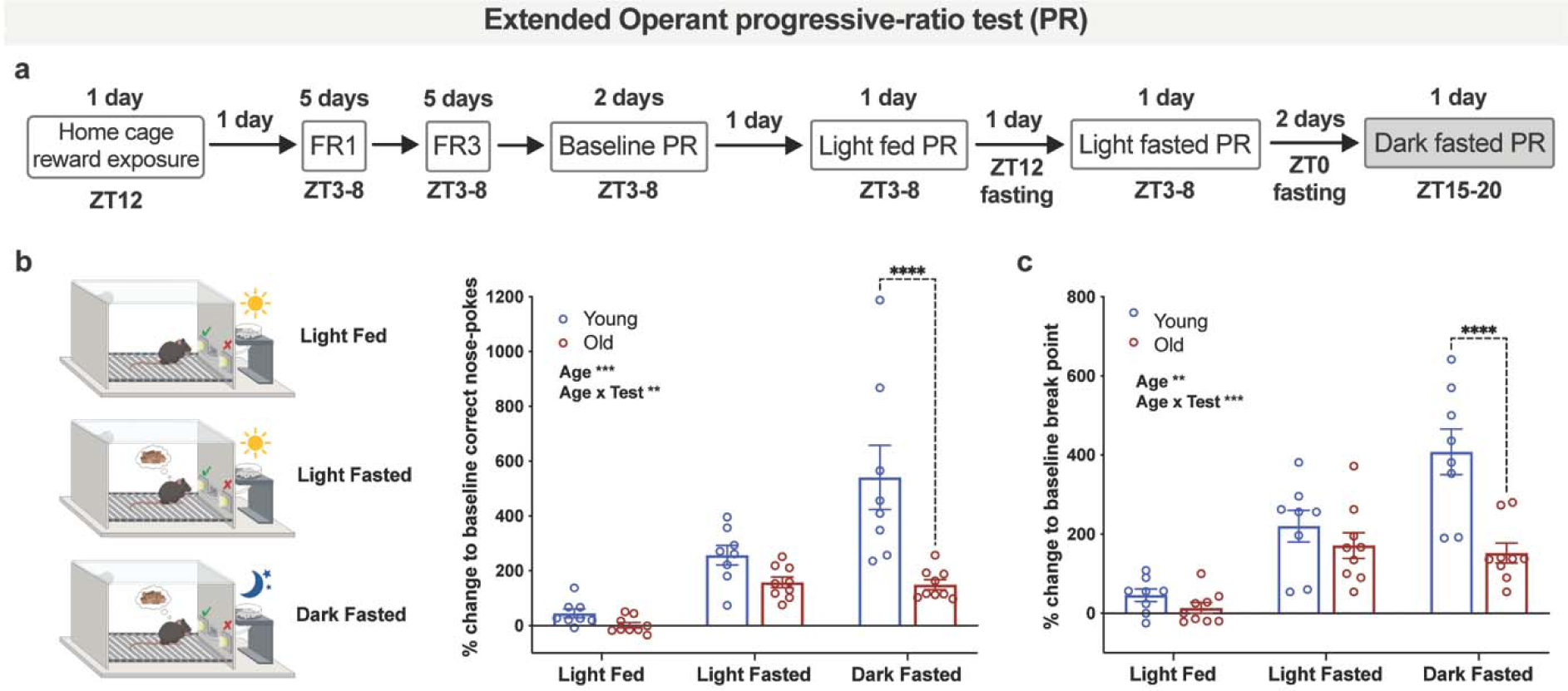
Aging leads to reduction of fasting-induced motivation for palatable food reward. Young (5 months) and old (19 months) mice were examined for motivation levels. (**a**) Calendar illustrating the extended operant progressive ratio test protocol. Correct nosepokes (**b**) and break points reached (**c**) normalized to individual baseline PR under different experimental conditions in the extended PR. Data were analyzed by 2-way ANOVA with *post-hoc* multiple comparisons. n=8-9. Error bars indicate SEM. Significant pair-wise comparisons and main effects of factors and interactions are labeled on the graphs: *p≤0.05; **p≤0.01; ***p≤0.001; ****p≤0.0001.

### BDNF mRNA and protein levels are decreased specifically in the ventral tegmental area (VTA) during aging

Extensive literature has defined the role of the mesolimbic dopamine (DA) system in regulating reward and motivation^16, 17^. Two major DA populations are identified in the ventral tegmental area (VTA) and the substantia nigra (SN) that locate adjacently in the ventral midbrain^18^. To determine the mechanism for this age-associated decline in motivation, we examined mRNA expression levels of characteristic functional genes in these two regions by collecting tissue samples from young and old brain sections with laser-capture microdissection and conducting quantitative RT-PCR. We found that expression of brain-derived neurotrophic factor (*Bdnf*) was significantly decreased in the aged VTA (Fig. 3a). Interestingly, genes related to DA turnover did not show any significant changes. The expression of *Gria1*, which encodes the GluA1 AMPA glutamate receptor subunit, tended to decrease in aged VTA (Fig. 3a). On the other hand, no genes examined in the SN were significantly altered during aging, except for a trend of increase in the expression of *Maoa*, which can be indicative for augmented DA metabolism and oxidative stress in this region (Fig. 3b)^19, 20^. We also confirmed this age-associated decrease in *Bdnf* expression by fluorescent *in situ* hybridization. Lower *Bdnf* signals were detected across eight consecutive bregma levels throughout the aged VTA, compared to the young VTA (Fig. 3c-d), whereas the expression of *Slc6a3*, a DA neuronal marker, did not show any significant differences between young and old mice (Extended Data Fig. 4a-b). Age-associated decreased *Bdnf* expression was present throughout the VTA in both *Slc6a3*-positive and *Slc6a3*-negative cells (Extended Data Fig. 4c-f), suggesting widespread downregulation of *Bdnf* throughout the VTA. Whereas aging selectively decreased *Bdnf* expression in the VTA, social isolation did not decrease *Bdnf* expression in the VTA (Extended Data Fig. 4g, left panel). On the other hand, social isolation tended to decrease *Bdnf* expression in the dorsal raphe nucleus (DRN) at least in young mice (Extended Data Fig. 4g, right panel). Given that the activity of DRN DA neurons is important to elicit a loneliness-like state and promote social preference^21,22^, a commonality between aging and social isolation appears to be decreased *Bdnf* expression, but in different areas of the brain. Following from the decrease in *Bdnf* mRNA, we further observed that matured BDNF (mBDNF) protein levels were moderately reduced in the aged VTA (Fig. 3 f-g), suggesting a potential involvement of *Bdnf* in age-associated reduced motivation.

**Fig. 3.**
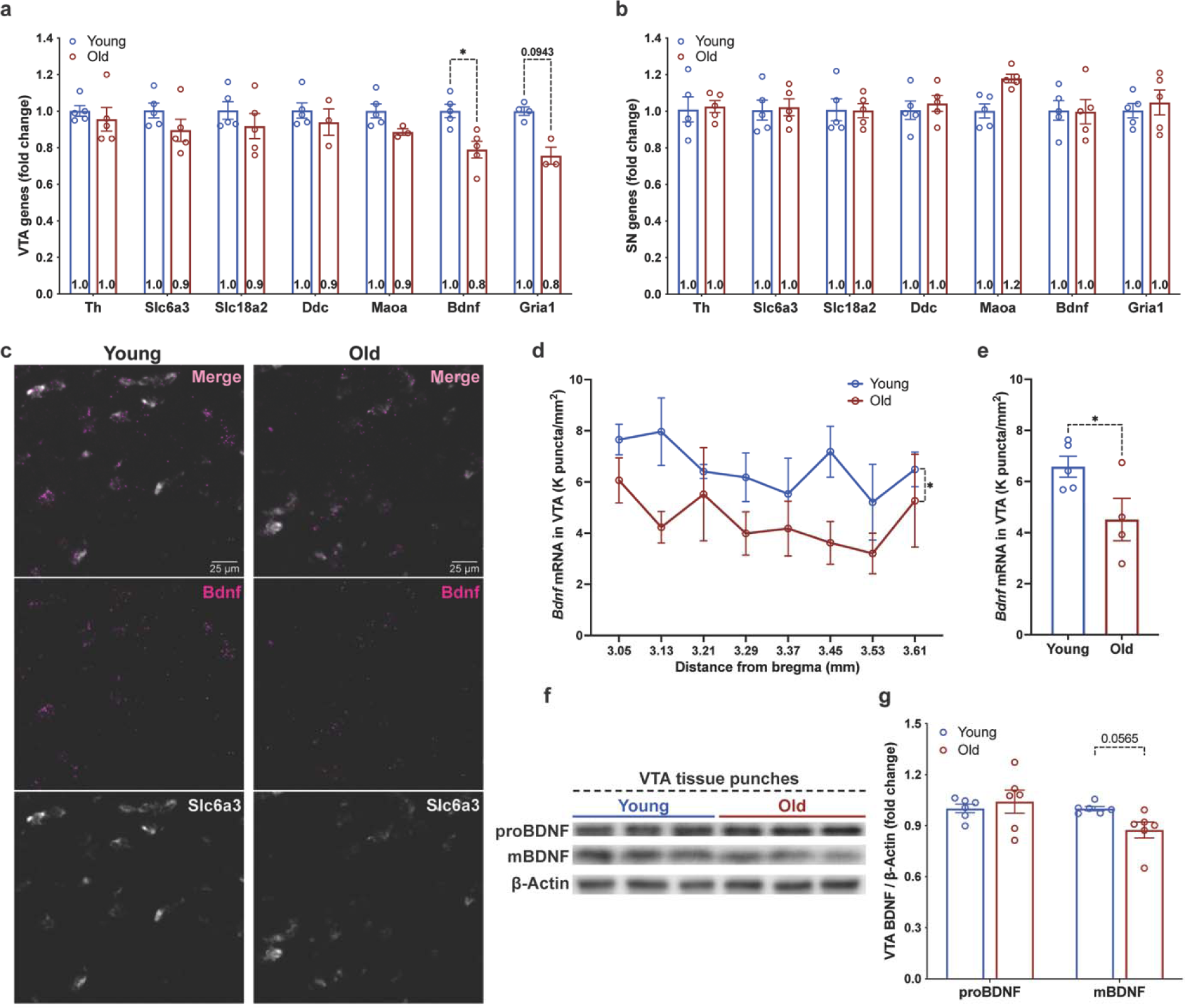
*Bdnf* gene expression is decreased in the ventral tegmental area (VTA) of aged mice. RT-qPCR analysis of laser-capture-microdissected VTA (**a**) and SN (**b**) samples from young (7 months) and old (21 months) mice. Representative images of FISH (RNAscope) analysis of *Bdnf* in VTA, using dopamine active transporter (DAT/*Slc6a3*) mRNA as a marker for DA neurons in young (7 month) vs. old mice (22 months) are shown in (**c**) and quantified as the density of *Bdnf* mRNA signal puncta in the VTA region (**d**, **e**) across 8 bregma levels (80μm interval from −3.05). Western blots (**f**) and quantifications (**g**) of the precursor and matured forms of the BDNF protein (proBDNF and mBDNF, respectively) normalized to β-actin signals in punched VTA of young (4 months) vs. old (23 months) mice. *Gria1*: glutamate ionotropic receptor AMPA type subunit 1; *Slc6a3*: solute carrier family 6 member 3. Data were analyzed by 2-way ANOVA with *post-hoc* multiple comparisons (**a**, **b**, **d**, **g**) and student’s t-test (**e**). n= 3-6. Error bars indicate SEM. Significant pair-wise comparisons and main effects of factors and interactions are labeled on the graphs: *p≤0.05; **p≤0.01; ***p≤0.001; ****p≤0.0001.

### VTA-specific *Bdnf* knockdown in young mice recapitulates the reduced motivation observed in aged mice

Following the observation that VTA *Bdnf* was decreased in aged mice, we attempted to restore VTA *Bdnf* expression in old mice to determine if their motivation could be rescued. To do so, we used a *Bdnf*-expressing lentivirus. This approach yielded significant BDNF protein overexpression in primary mesencephalon neuronal culture, but it failed to increase BDNF protein levels *in vivo* so that PR performances were not improved in old mice (Extended Data Fig. 5). We suspect that this could be due to certain auto-regulatory mechanisms that suppress *Bdnf* transcription and/or translation in the VTA neurons in response to overexpression^23–26^. It is also possible that the lentiviral transgene was silenced in the host cells over time^27–29^. We next sought to determine whether decreasing VTA *Bdnf* expression in young mice can recapitulate the motivational dysfunction observed in aged mice. To do so, we knocked down *Bdnf* expression in the VTA by stereotactically injecting a *Bdnf* shRNA-expressing lentivirus into the VTA of young mice (Fig. 4a). *Bdnf* mRNA levels were reduced by ∼30% (Fig. 4b-c), and the mBDNF protein levels were also reduced to a similar extent to aged mice in the VTA, but not in the downstream nucleus accumbens (NAc) (Fig. 4d-f). Two months after surgery, the VTA-specific *Bdnf*-knockdown (VTA-*Bdnf*-KD) mice were tested for the extended PR paradigm with food deprivation during light and dark times. Importantly, VTA-*Bdnf*-KD mice exhibited significant reduction in correct nose-pokes and break points with food deprivation during both light and dark times, successfully recapitulating motivational phenotypes observed in aged mice (Fig. 4g-h, and Extended Data Fig. 6a-b). On the other hand, VTA-*Bdnf*-KD mice did not alter behaviors in the VWR and OFT (Fig. 4i-l, and Extended Data Fig. 6c-d), supporting a selective role of *Bdnf* in the VTA in reward-seeking motivational behavior. Prior work has shown that VTA BDNF can play a role in regulating VTA dopamine neuron excitability^30, 31^. To determine whether the downregulation of VTA *Bdnf* in aged mice altered excitability in these VTA neurons, we examined the intrinsic neuronal electrical properties of both putative dopaminergic and non-dopaminergic neurons in young and aged mice. For these purposes, we operationally defined these neurons by the expression of hyperpolarization-activated cation current (I_h_)^32^ (Extended Data Fig. 7a-b). No significant differences were found between young and aged mice, suggesting that baseline neuronal excitability in the VTA does not drive the decreased motivation through aging (Extended Data Fig. 7c-f). Taken together, these results demonstrate that specific decreases in *Bdnf* expression in the aged VTA contribute, at least in part, to age-associated reduced motivation in mice.

**Fig. 4.**
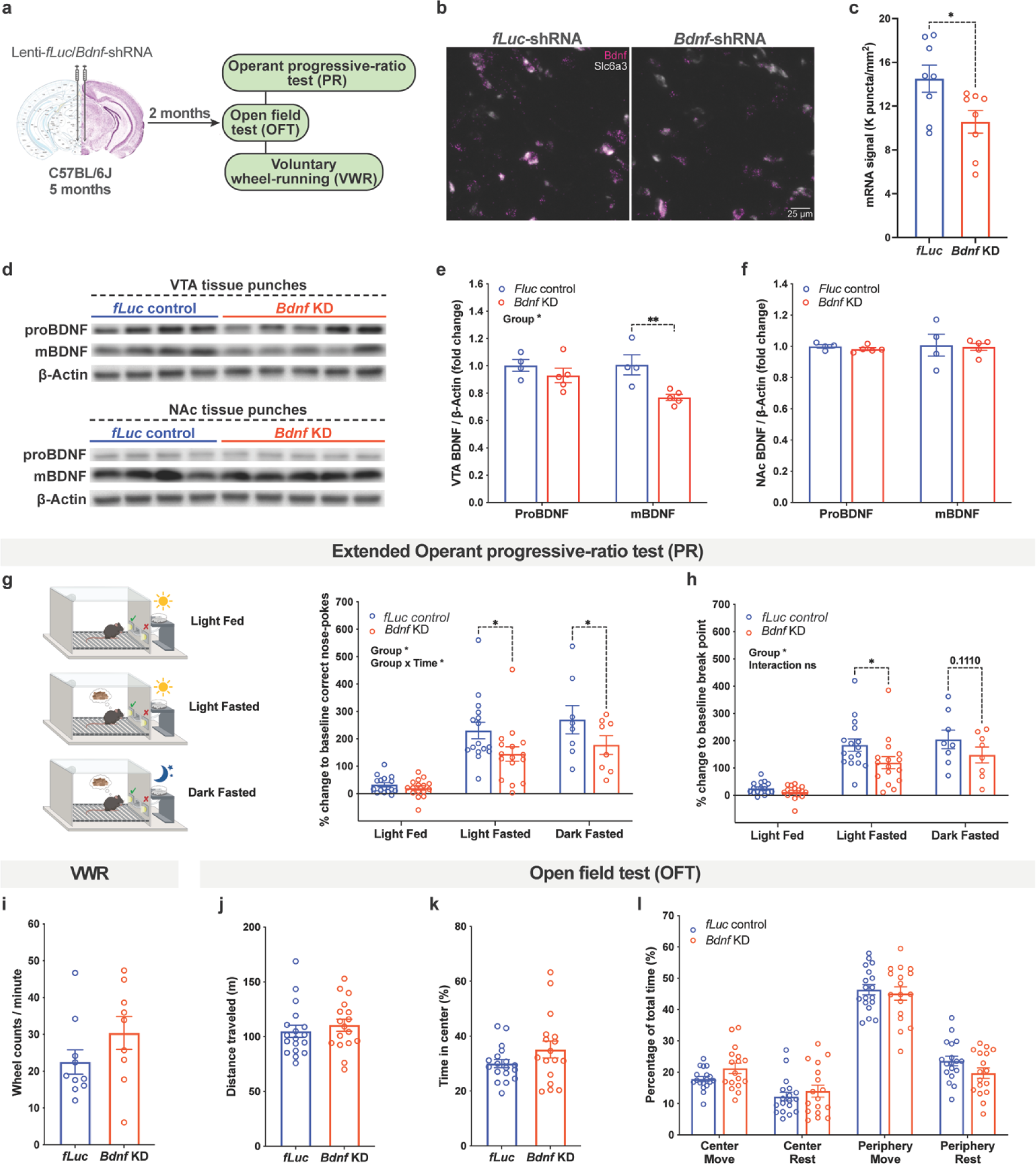
VTA *Bdnf* KD in young mice partially recapitulates the age-associated reduction in palatable food motivation, not affecting other age-related phenotypes. (a) Calendar and schematics illustrating the experimental design. (**b**) Representative FISH images of *Bdnf* and Slc6a3 in VTA of *fLuc* vs *Bdnf*-shRNA virus injected mice after behavior tests, (**c**) quantified as the density of *Bdnf* mRNA signal puncta in infected regions (KD efficiency 27%). Western blots (**d**) and quantifications (**e, f**) of proBDNF and mBDNF levels normalized to β-Actin signals in punched VTA and NAc of *fLuc* vs *Bdnf*-shRNA virus injected mice. Correct nosepokes (**g**) and break points reached (**h**) normalized to individual baseline PR under different experimental conditions in the extended PR. (**i**) Average VWR activity during the dark phase (ZT12-0). Total distance traveled (**j**), percentage of time spent in the center area (**k**) and percentage of time spent moving or resting in the center and peripheral areas (**l**) in the open field test. Data were analyzed by 2-way ANOVA with *post-hoc* multiple comparisons (**e**, **f**, **g**, **h**,), student’s t-test (**c**, **i**, **j**, **k**), and Holm-Šídák multiple comparison test (**l**). n=4-18. Error bars indicate SEM. Significant pair-wise comparisons and main effects of factors and interactions are labeled on the graphs: *p≤0.05; **p≤0.01; ***p≤0.001; ****p≤0.0001.

## Discussion

Here we demonstrated that both social isolation and aging significantly decreased motivation in mice along with the induction of certain anxiety- or depression-related behaviors. Aging is a strong contributor to reduced motivation, and this may explain why aged mice respond substantially less to social isolation compared to young mice. In this regard, we found that *Bdnf* expression was significantly reduced throughout the aged VTA, in both DA and non-DA cells. Remarkably, the VTA-specific knockdown of *Bdnf* in young mice recapitulated the reduced motivation observed in old mice, suggesting that this VTA-specific reduction in *Bdnf* contributed to the impaired reward-seeking motivation in aged mice. Whereas VTA BDNF levels have been implicated in the development and progression of drug-induced reward processing or addiction^33, 34^, our findings suggest that BDNF also plays an important role in maintaining motivated states in a healthy, physiological conditions, which impair with age. However, given that VTA-*Bdnf*-KD mice did not alter voluntary wheel-running and open field activities, the effect of age-associated BDNF reduction is limited to palatable reward-seeking motivation. Furthermore, because neuronal excitability in the VTA shows no significant changes during aging, it is possible that endogenously evoked activity and synaptic transmission in this particular reward-seeking circuit may be affected by age-associated BDNF reduction.

Importantly, while the high frequency of depression in Parkinson’s disease appears dependent on decreased DA innervation of the limbic system^35, 36^, the age-associated decreases we observed in VTA *Bdnf* expression occurred in both DA and non-DA cells, suggesting that this is a distinct mechanism for age-associated decreased motivation. Interestingly, it has been reported that leptin receptor-expressing neurons in the lateral hypothalamus (LH) mainly project to non-DA VTA neurons and drive PR performance through the activation of these VTA neurons ^37^. We previously showed that specific neuronal populations in the dorsomedial hypothalamus and LH play critical roles in counteracting age-associated functional decline and determining longevity in mice ^38–40^. Therefore, it will be of great interest to determine whether these neuronal populations that fundamentally control aging and longevity also regulate motivation through *Bdnf*-expressing VTA neurons. Because the LH shows significant reduction in its NAD^+^ levels during aging and may rely on extracellular nicotinamide phosphoribosyltransferase (eNAMPT), an important NAD^+^ biosynthetic enzyme counteracting aging, for its NAD^+^ biosynthesis^41–43^, it will be important to examine whether eNAMPT supplementation could restore motivation in aged mice.

In humans, it has been reported that BDNF in the brain decreases with age^44–47^. In particular, Buchman *et al.* reported that in more than 500 older adults, high *BDNF* expression levels in the brain were associated with slower rates of cognitive decline^45^. Additionally, Oh *et al.* showed that age-associated *BDNF* decline was assocaited with the downregulation of excitatory and inhibitory synaptic markers, implicating that *BDNF* reduction may affect synaptic transmission in the aged brain^44^. Although previous studies did not assess the connection between *BDNF* and motivation, it is likely that age-associated *BDNF* decline in the brain could cause reduced motivation in humans. Interestingly, young adult human subjects (age 18-29) perceive the highest levels of isolation during the pandemic of COVID-19, compared to old subjects (age 50-69), and overall life satisfaction is significantly lower for people who perceived greater social isolation^48^. This trend is qualitatively similar to our results in the PR test, showing that aged mice are less responsive to social isolation, compared to young mice, most likely due to already reduced levels of motivation. Therefore, our results in this study could provide an important insight into age-associated alterations in mental states in humans.

In conclusion, the present study advances our understanding of the mechanism of age-associated motivational changes and the physiological importance of Bdnf in the VTA. It also stimulates development of interventions to keep our motivation levels high and healthy during aging.

## Methods

### Animals

All animal procedures were approved by the Washington University Institutional Animal Care and Use Committee and were in accordance with National Institutes of Health guidelines. Unless otherwise noted, all mice were housed in groups of 2-5, fed *ad libitum* on a standard chow diet (PicoLab Rodent Diet 20-5053; LabDiet, St. Louis, MO) and autoclaved water in a temperature/humidity-controlled holding room within a barrier facility, maintained on a 12:12-hour light/dark cycle. Food, beddings (Aspen Chip, Northeastern Products Corp.), and nesting material (Nestlets, Ancare) were changed once per week; water bottles were changed as needed. Mice were labeled on their tails with sharpie markers for easy identification and monitored constantly for their health status. All handling, cage changing, measuring procedures, and behavior tests were conducted by the same experimenters throughout the study. Only male mice were used.

All aged mice and the corresponding young groups were obtained from the NIH aged rodent colony (C57BL/6JN) or our in-house C57BL/6J colony. Due to the limited availability of aged mice, this study only used male mice to limit the number of potential biological variables to remain sufficiently powered. All group-housed cages were housed only with original cage mates from weaning. Mice used for *Bdnf* knock-down analysis were purchased from Jackson Laboratories (C57BL/6J) at 12-week-old and acclimated to the animal facility for ∼2 months before surgeries; upon arrival, mice were randomly assigned into cages of 4/5 and maintained until sacrifice.

### Social isolation procedure

All mice were originally group-housed in cages of 3-5 cage mates. After acclimation to the holding rooms, cages containing around half of the animals were separated into individual cages and maintained isolated until sacrifice; the rest of the control cages remained unchanged across the study. All cages within a certain cohort were placed in proximity on the same rack to control for traffic, noise levels, and light exposure.

### Behavior tests

All behavioral tests were performed within a sound-attenuated room maintained at 20-26°C after at least one week of acclimation to the local holding room. All tests were performed by female experimenters. Unless otherwise noted, all mice in the same cohort underwent the same test/procedure on the same day(s) from ZT3 to ZT9; if a test/procedure cannot be finished within one day, the cohort will be divided into sub cohorts by counterbalancing grouping, home cages, and other factors. For all experiments performed outside of the holding room, mice were brought into the corresponding behavior space to habituate for at least 20 minutes before any test/procedure started. All behavioral apparatuses were cleaned with 2% chlorhexidine in between subjects.

#### Operant Progressive-Ratio Test

To expose the animals to the reward, mice fed *ad libitum* were given 3 g/mouse of the reward sugar pellet (Dustless Precision Pellet, F0071, Bio-Serv) on the bedding of their home cages when cages were changed at ZT12. Consumption of the pellets was confirmed the next day. One day after the exposure, mice were gently placed in a mouse operant-conditioning box (15.24×13.34×12.7 cm) within a sound attenuating cubicle (ENV-307A & ENV-020M, Med Associates Inc., Fairfax, VT). The box was equipped with a stainless steel grid floor, two nose holes (active and inactive) flanking a trough pellet receptacle (food tray) connected to a pellet dispenser on the bottom of the right-hand side wall; a house light was installed at the top center of the left-hand side wall. Pokes into the ports are monitored by infrared light beam sensors. Upon starting of a fixed-ratio 1 (FR1) session, both nose holes present a cue light; one nosepoke into the active hole results in a sugar pellet (20 mg) dispensed to the food tray, along with cue lights turned off and house light turned on for 2 s, during which no action triggers event (time-out period); poking the inactive hole has no consequence. Each session lasted 60 minutes. In between trials, boxes were cleaned with 2% chlorhexidine, and leftover pellets in the dispenser were replaced with fresh and intact ones. All trials took place from ZT3 to ZT8, unless otherwise specified, and each animal was subjected to the sessions at the same time across days. In FR1, mice were trained to discriminate the active hole from the inactive one and receive rewards from the food tray. Following 5 consecutive days of FR1 sessions, mice were moved on to another 5 consecutive days of 60-minute fixed-ratio 3 (FR3) sessions, in which three active nosepokes results in one reward delivery and 20 s time-out period. Mice were conditioned to repeatedly perform the correct operant behavior until receipt of the reward during FR3. After completion of the training, motivation levels of the mice were assessed using a progressive-ratio (PR) schedule of reinforcement following the equation: response ratio = 5*e^reward^ ^number/^*^5^ – 5 (rounded to the nearest integer), resulting in an exponential increase of the number of correct pokes required for each subsequent reward: 1, 2, 4, 7, 9, 12, 15, 20, 25, 32, 40, 50, 62, 77, 95, 118, 145, 178, 219, 268, 328, 402…^49^. All events were programed, and actions were recorded using MED-PC software (Med Associates) controlled by a PC. The number of active nosepokes counted toward rewards during the 60-minute session was calculated as the readout of motivation; A break point (BP) was calculated by taking the bigger number of the last completed ratio versus the number of correct nosepokes after the last reward^13^. For the extended PR paradigm, following FR1 and FR3 training, two PR sessions were conducted on consecutive days, and the averaged readouts served as the PR baseline. Mice then underwent a break day with no session, followed by a third PR test (light fed) under the same conditions (*ad libitum* in the light time); after another break day and overnight fasting from ZT12, a fourth PR test was conducted (light fasted); lastly, two days after, the mice were fasted from ZT0, and a fifth PR test under fasting was conducted during ZT15-20 under dim red lighting (∼10 Lux).

#### Sucrose Preference Test

Mice were habituated to the sipper tubes in their original home cages by replacing the drinking bottle with a sipper tube for 4 days; daily water consumption was confirmed to be > 2 mL/mouse/day by the last day of the habituation. The sipper tube was home-constructed with a cut 25-mL plastic serological pipettes and a stainless steel pin valve (Lixit) sealed together with heat shrink tubing. Mice were then all separated into individual cages (if not already separated as part of the social isolation procedure), given *ad libitum* regular chow and two identical sipper tubes positioned side-by-side next to the chow, both containing drinking water. Baseline water consumption was measured by recording volume left in each sipper tube daily for 4 days. After that, water in one of the tubes was replaced with 2% sucrose solution. Sucrose solution and water consumed per day were recorded for another 4 days, and the percentage of sucrose solution over total liquid intake was calculated to represent sucrose preference. Tube positions (left vs right) were switched daily after volume checking across the entire test. Mice were housed back with their original cage mates after the procedure was finished.

#### Elevated-Plus Maze Test

The EPM was performed under dim warm lighting stabilized at ∼5-6 lux. The apparatus was a plus-shaped platform comprised of two open arms (25×5×0.5 cm) across from each other and perpendicular to two closed arms (25×5×16 cm) with a center platform (5×5×0.5 cm), elevated to be 50 cm above the floor (60140, Stoelting Co). After habituating to the room, each mouse was individually placed onto the apparatus from one of the open arms and allowed to freely explore. Activity was video recorded using a CCD camera for 20 minutes and videos analyzed using Ethovision XT 13 (Noldus Information Technology, Wageningen, The Netherlands). Behavior tracking after the mice made the first entry into the center area was included in the analysis.

#### One-hour Locomotor Activity and Open Field Test

The 1-hour LA/OFT was conducted in the Animal Behavior Core at Washington University School of Medicine in St. Louis. The test was performed under standard room lighting stabilized at ∼150 lux The apparatus consisted of a transparent (40.6×40.6×38.1 cm) polystyrene enclosure, surrounded by a frame containing a 16×16 matrix of photocell pairs. Output was fed to an online computer and analyzed by the system software (Hamilton-Kinder, LLC, Poway, CA). A center area was defined as a 9×9 grid within the 15×15 grid in this chamber, with the peripheral area extending to the sides of the chamber as the outermost boundary. After habituating to the room, each mouse was individually placed in the center of the apparatus and allowed to freely explore for one hour. The OFT for VTA-*Bdnf*-KD mice was performed under dim warm lighting stabilized at ∼5-6 lux. The apparatus consisted of an open-top opaque square enclosure (50×50×31 cm). After habituating to the room, each mouse was individually placed at the center of the apparatus and allowed to freely explore. Activity was video recorded using a Google Pixel 3 XL smartphone for 30 minutes and videos analyzed using Ethovision XT 13. The center was defined as a square comprised of 36% the total area of the OFT (i.e., each side was 0.6 that of the total arena).

#### Hole-board olfactory preference test

The HOP was conducted in the Animal Behavior Core at Washington University School of Medicine in St. Louis. The test was performed under standard room lighting stabilized at ∼150 lux. The apparatus consisted of a transparent (40.6×40.6×38.1 cm) polystyrene enclosure equipped with a computerized hole-board apparatus of the same size, which contains 4 corner and 4 side holes, with a side hole being equidistant between the corner holes (Learning Holeboard; MotorMonitor, Kinder Scientific, LLC, Poway, CA). This task consisted of two consecutive days. On day 1, mice were put into the apparatus for 20 min to habituate. On day 2, mice were presented with one of 8 options for configurating of corner holes containing either female urine, vanilla extract, distilled water, or empty. Pairs of photocells are equipped within each hole (27 mm diameter) and were used to quantify the frequency and duration of pokes. Specifically, a mouse was required to place its head into a hole at a depth of at least 35 mm and break a photobeam for the response to be registered as a hole poke. All odorant-containing cups were cleaned with mild soap and water, and the entire apparatus was cleaned with a 2% chlorohexidine diacetate solution to remove the odor between trials.

#### Social Interaction Test

The test was performed in recording closets under lighting stabilized at ∼ 10 lux. The apparatus consisted of a homemade black opaque two-chamber box (52×26×26 cm) divided in half by blockers on the sides, leaving the center open (W10×H26 cm). A metal, mesh pencil cup (10H×8.5 radius cm) was invertedly placed in the center of each chamber, with extra weight and a cone cup attached on top to prevent animals from moving or climbing on them. The mesh allows for exchange of sensory information and prevents direct physical interactions between animals inside and outside of the cup. The floor of the box was covered by a thin layer of bedding material. After habituating to the room, to assess baseline activities, each subject mouse was individually placed in the center of the box and allowed to freely explore for 10 min, followed by 30 min back to their home cages. To test for sociability, a sex- and age-matched stranger conspecific was placed under one of the cups in a counterbalanced and randomized manner, and the subject mouse was then reintroduced to the same box and allowed to freely explore for another 10 min. Bedding in the boxes was replaced with fresh one in-between subjects. Activity was video-recorded using a CCD camera, and video recordings were analyzed using Ethovision XT 13. Activities of the mice in either chamber during the baseline and the test phases were analyzed.

### Voluntary wheel-running activity

After concluding prior behavior test(s), mice were transferred to the VWR facilities and housed individually in a Tecniplast polycarbonate cage (33.2×15×13 cm) (PT2-MCR, Phenome Technologies) with a vertical running wheel (11 cm inside diameter, 5.4 cm wide), stainless steel wire top, and a filter cage cover (Actimetrics). Wheel-running behavior was recorded and analyzed using the ClockLab software (https://actimetrics.com/products/clocklab/). Mice were allowed to habituate to the wheel cages for at least a week and acquire stable running activity before data was collected.

### Laser-captured microdissection

Mice were sacrificed by CO_2_. Brains were removed, segmented into separate parts in a brain matrix, and immediately embedded in OCT and snap frozen by powered dry ice. Brain samples were stored at −80°C until sliced into 30 μm coronal sections in a cryostat at −14°C. Sections were thaw-mounted onto PEN-membrane slides (#11505158, Leica Microsystems, Buffalo Grove, IL, USA) and kept frozen until the next step. After collecting all sections containing brain regions of interest, slides were hydrated sequentially in 100%, 95%, 75%, and 50% ethanol for 30 sec each. The hydrated slides were stained with 0.5% Cresyl Violet (Sigma-Aldrich) for 2 minutes, and dehydrated with 50%, 75%, 95%, and 2 rounds of 100% ethanol for 30s each. The dehydrated slides were then incubated in xylene twice for 1 minute each. After being air-dried, VTA, SN, and DRN regions were microdissected using the Leica LMD 6000 laser-microdissection system, following the Franklin and Paxinos mouse brain atlas. Laser-cut tissue samples were collected in the extraction buffer of the Arcturus PicoPure RNA Isolation Kit (12204-01, Applied Biosystems), and stored at −80°C until RNA was isolated following the manufacturer’s protocol and DNase (79254, Qiagen) treatment.

### Quantitative real-time RT-PCR

Concentrations of RNA samples were measured by NanoDrop and stored at −80°C until being reverse-transcribed into cDNA with the High Capacity cDNA Reverse Transcription kit (#4368814, Thermo Fisher). qRT-PCR was conducted with the TaqMan^TM^ Fast Universal PCR Master mix (4352042, Applied Biosystems) and TaqMan primers for each gene (*Gapdh*: Mm99999915_g1, *Th*: Mm00447557_m1, *Slc6a3*: Mm00438388_m1, *Slc18a2*: Mm00553058_m1, *Ddc*: Mm00516688_m1, *Maoa*: Mm00558004_m1, *Bdnf*: Mm04230607_s1 and 4331348_APNKZZR, a customized primer for the overexpressed *Bdnf* mRNA: ATAGCAAAAAGAGAATTGGCTGGCGATTCATAAGGATAGACACTTCCTGTGTATGTACACTGACCATTAAAAGGGGAAGA, *Gria1*: Mm00433753_m1) in the StepOnePlus^TM^ Real-Time PCR System (Applied Biosystems). Relative expression levels were calculated for each gene by normalizing to *Gapdh* CT values and then to the mean of the control/reference group.

### RNAscope Fluorescent *In Situ* Hybridization (FISH)

Mice were transcardinally perfused first with 0.01M phosphate-buffered saline (PBS) and then the fixative: 2% paraformaldehyde (PFA) and 0.2% picric acid in 0.01M PBS, pH 7.4. Brains were removed and post-fixed in the fixative at 4°C overnight, followed by cryoprotection in 20% sucrose in 0.01M PBS at 4°C for at least 24 hours and snap freezing by powered dry ice. Brains were stored at −80°C until sliced into 20 μm coronal sections containing the VTA region in a cryostat at −20°C. Sections were thaw-mounted onto Superfrost^TM^ Plus Microscope Slides (12-550-15, Fisher Scientific) and kept frozen until use. FISH was performed using RNAscope Multiplex Fluorescent Reagent Kit v2 [Advanced Cell Diagnostics, Inc. (ACD)] as per the manufacturer’s protocol. Briefly, slides washed with 1x DPBS were fixed in pre-chilled 4% PFA in 1x DPBS at 4°C for 15 minutes. Sections were then dehydrated through 50%, 70%, and two rounds of 100% ethanol for 5 minutes each, followed by air-drying. Then, the sections were treated with RNAscope® Hydrogen Peroxide for 10 minutes at room temperature. A hydrophobic barrier was created around the sections using ImmEdge^TM^ Hydrophobic Barrier PAP Pen (H-4000, Vector Laboratories) and allowed to air-dry. The slides were then incubated with Protease III for 30 minutes and hybridized with ACD probes for target genes (Mm-*Bdnf*-CDS, #457761, probe region 662-1403; Mm-*Slc6a3*-C3, 315441-C3, probe region 1486-2525) for 2 hours at 40°C using a HybEZ^TM^ oven. The signals were amplified by subsequent incubations with Amp-1, Amp-2 and Amp-3 reagents for 30, 30, and 15 minutes, and then with the HRP reagents, depending on the channels used, in combination with the TSA Plus (NEL760001KT, PerkinElmer) and Opal (FP1487001KT, Akoya) fluorophores at 1:1000 to 1:500 dilutions. Sections were washed with 1x RNAscope Washing Buffer in-between every incubation. Slides were counterstained with DAPI, and coverslips were mounted with FluorSave™ Reagent (#345789, Calbiochem). Images were obtained using fluorescent microscopy (DMi8, Leica Microsystems) under 20x magnification. Fluorescent signals were quantified using a customized ImageJ (NIH) pipeline. Briefly, after tiff images were exported, a circular ROI (radius = 200 μm) was selected at the center of the VTA region, referring to the shape covered by the *Slc6a3* signal; after subtracting background from all channels, the *Slc6a3* channel underwent Gaussian Blur, binary conversion, and shape growth to approximate the area covered by the dopaminergic neurons; the number of the *Bdnf* puncta was quantified using the Find Maxima function, and the identified points were overlayed on the *Slc6a3* area to count the *Bdnf* signals that are expressed in the dopaminergic neurons. For the *Bdnf* knock-down experiment, signals were quantified where the virus caused infection in the VTA.

### Brain punching

Mice were transcardinally perfused with 0.01M PBS, and brains were removed and snap frozen by powered dry ice. Brain samples were stored at −80°C until sliced into 200um coronal sections from anterior to posterior of the brain in a cryostat at −8°C. Section containing the nucleus accumbens (NAc) and VTA regions were identified by bregma landmarks and punched to acquire these regions using a Brain Punch Set (#57401, Stoelting). The NAc region were collected by the 1.19 mm punch from one side of 4 consecutive brain sections (Bregma around 1.5 to 0.7); the VTA region were collected by the 0.74 mm punch from both sides of 5 consecutive brain sections (Bregma around −2.8 to −3.8). To proceed to western blotting, each punched sample was homogenized in 60 μL 1x sample buffer containing 10% 2-Mercaptoethanol by passing through a syringe with 28-gauge needle (102-SN1C2805P, McKesson) for 5 times. The homogenate was then centrifuged at 16,000 xg for 2 minutes, and the clear upper layer of 55 μL was carefully collected into a new tube.

### Western blotting

Protein samples prepared in 1x sample buffer containing 10% 2-Mercaptoethanol were incubated at 95°C for 10 minutes and allowed to cool down. 14 μL of each sample was electrophoresed on a 4-15% Mini-PROTEAN^®^ TGX™ Precast Protein Gels (#4561086, BioRAD) in a Tris-Glycine running buffer, at 80-100V. The separated protein samples were then transferred to a Immobilon^®^-P PVDF Membrane (IPVH00010, Millipore) in iced transfer buffer that contains 0.1% SDS and 10% methanol, at 0.2A for 50 minutes in a cold room. After being washed in 1x Tris-buffered saline with 0.1% Tween-20 (TBS-T) and blocked in 5% w/v skim milk/TBS-T for 1 hour at room temperature (RT), the membrane was then incubated with rabbit anti-BDNF (1:1000, ab108319, Abcam) or rabbit anti-β-actin (1:1000; PA1-183, Thermo Fisher) in 5% skim milk overnight at 4°C. After the membranes were washed in TBS-T for 3 times, they were incubated with HRP conjugated anti-rabbit IgG antibody (1:10,000, NA9340, Cytiva) in 5% skim milk for 1 hour at RT. After washing the membrane in TBS-T for 3 times, chemiluminescence signals were developed using Amersham ECL reagents (RPN2235 or RPN2232, Cytiva) and visualized using the [647SP, no light] setting of the ChemiDoc MP Imaging System (BioRAD) with exposure times of 1-20 minutes. When need to be reblotted, membranes were stripped with a stripping buffer (#21059, Thermo Fisher) for 15-30 minutes at RT. All incubations and washes were done with constant rocking. Signal intensity of the appropriate bands was quantified using ImageJ (NIH).

### Virus vector construction

To generate the shRNA expressing lentiviral constructs, 56-bp double-stranded oligonucleotides, each of which contains a sense target sequence, a microRNA based loop sequence (CTTCCTGTCA), an antisense sequence, and a 4-nucleotide sequence complimentary to the vector cutting site of the restriction enzyme Aarl at both ends, were generated for *Bdnf* and firefly luciferase (*fLuc*) and cloned into the U6-PGK-eGFP vector, provided by the Hope Center Viral Vectors Core at Washington University School of Medicine in St. Louis. Three shRNA targets on the *Bdnf* coding sequence were cloned and tested in parallel, and the one that gave the highest knock-down efficiency *in vitro* was selected for *in vivo* experiments. The sense sequence used for *Bdnf* was 5’-ACTATCCATTCTGGTTGATAA-3’ and 5’- TACGCGGAATACTTCGAAATG-3’ for *fLuc*.

Gibson Assembly designed using the NEBuilder™ website (https://nebuilder.neb.com) was used to generate the *Bdnf* overexpressing lentiviral constructs. The *Bdnf* cDNA without stop codon was cloned from total cDNA extracted from a wildtype mouse hippocampus sample using the primer set: 5’-CCGTTACTAGTGGATCCACCGGTGCCACCATGACCATCCTTTTC-3’ and 5’-CGGAGCCTCTTCCCCTTTTAATGGTCAG-3’. A P2A-eGFP sequence was cloned from the AAV-CamKIIa-hWFS1-P2A-eGFP vector, provided by the Hope Center Viral Vectors Core, using the primer set: 5’-GGGGAAGAGGCTCCGGAGCCACGAAC-3’ and 5’- AATTCCGCGGGCCCGTCGACTTACTTGTACAGCTCGTCCATGC-3’. The two segments were then assembled into the LV-SYN-eGFP vector, provided by the Hope Center Viral Vectors Core, using the Gibson Assembly® Master Mix (E2611, NEB). The original LV-SYN-eGFP vector was used as the control. After confirming the new constructs by sanger sequencing provided by GENEWIZ, high concentrations of vector DNA samples were produced using ZymoPURE II Plasmid Maxiprep Kit (D4203, ZYMO), following the manufacturer’s protocol. High-titer lentiviruses were produced by the Hope Center Viral Vectors Core. Briefly, HEK293T cells were co-transfected with the shRNA-expressing vectors and three packaging vectors (pMD-Lg/pRRE, pCMV-G, and RSV-REV) by using polyethylenimine (PEI) reagent. Six hours after transfection, the medium was replaced with the complete medium containing 6 mM sodium butyrate. Culture supernatant was collected 42 hours after transfection. The supernatant was passed through a 0.2 μm filter, concentrated by ultracentrifugation through a 20% sucrose cushion, and stored at −80°C until use. Virus titer was determined by transducing HT1080 cells and assaying for vector sequence using qPCR. Virus with a titer of 1.1-1.7 x 10^9^ IU/mL was used for stereotaxic injection.

### Primary mesencephalon culture

Primary mesencephalon culture was prepared as previously described with modifications^50, 51^. Briefly, embryonic-day 12.5 (E12.5) C57BL/6JN mouse embryos were removed from timed pregnant females and placed in washing medium containing DMEM-High glucose (12800-017, Gibco) with antibiotics (15240-062, Gibco) on ice. The embryos were dissected to isolate the midbrain region, which was further trimmed to obtain the ventral mesencephalon tissue. The collected tissue was stored in and rinsed with iced washing medium for 2-3 times in a 15 mL conical tube before being digested in 1 mL (or 0.1 mL/embryo, which ever that is larger) 0.25% trypsin (25200-056, Gibco) with 0.5 mg/mL DNase (DN25-100MG, Sigma-Aldrich) in a 37°C water bath for 15 minutes. After gently swirling the mixture every 5 minutes, enzymatic activity was neutralized by adding an equal volume of plating medium comprised of DMEM/F12 (1:1) (11330-032, Gibco), 0.1 mg/mL glucose (25-037-CI, Corning), B27 supplement (17504-04, Gibco), and antibiotics (15240-062, Gibco). After mixing the content 3-5 times and waiting until tissue chunks settled down to the bottom of the tube, the supernatant was carefully removed with a pipette and slowly added 0.7-1.0 mL plating medium. Tissue was dissociated by gently pipetting the tissue ∼15 times with a flamed glass pasture pipette. The tube was kept undisturbed for 1-2 minutes to allow undissociated tissue debris to settle down, and the upper phase containing dissociated cells was collected to a new tube. The cell density was calculated using a hemocytometer (#1492, Hausser Scientific), and then the cell suspension was diluted with the plating medium to 1×10^6^ cells/mL. The cells were seeded at a density of 120,000 cells/cm^2^ on a poly-D-lysine (P6407, Sigma-Aldrich; 200 mg/mL)-coated culture plate. After 4-6 hours, the medium was replaced to the complete culture medium comprised of Neurobasal medium (21103-049, Gibco), GlutaMAX Supplement (35050-51, Gibco), B27 supplement (17504-04, Gibco), and antibiotics (15240-062, Gibco). To maintain the culture, half of the medium was changed every 3 days. Viruses were added to seeded wells at 1:200 ratio by volume after 7-8 days *in vitro* (DIV). Cells were harvested 72 hours after infection. RNA samples were extracted using the PureLink RNA Mini Kit (#12183025, Thermo Fisher) with DNase (#79254, Qiagen). Protein extracts were prepared using 1x sample buffer containing 10% 2-mercaptoethanol (0.1 mL/cm^2^) and centrifuged at 16,000 xg for 2 minutes. Supernatants were collected into new tubes and subjected to Western blotting.

### Stereotaxic injection

Mice were anaesthetized in an induction chamber (2-4% isoflurane) and placed in a stereotaxic frame (Model 942, Kopf Instruments) where they were maintained under 1-2% isoflurane on a 37°C heating pad throughout the procedure. Following craniotomy under aseptic techniques, all injections were done using a Hamilton Neuros Syringe (65460-03, Hamilton) driven by a Micro4 microsyringe pump controller (World Precision Instruments, Sarasota, FL), with the beveled opening facing posterior. The syringe was slowly inserted into the brain tissue and dwelled in place for 3 minutes before the injection started; after injection, the syringe was dwelled for another 1 min/100 nL virus injected and then slowly withdrawn. The skin incision was closed with 6-0 nylon sutures. For the VTA *Bdnf* knock-down experiments, mice were injected bilaterally with 1 μL of either lenti-shRNA-*Bdnf*-eGFP or lenti-shRNA-*fLuc*-eGFP at the rate of 200 nL/min into the VTA (coordinates relative to Bregma: AP −3.05, ML ±0.3, DV −4.6). For the VTA *Bdnf* overexpression experiments, mice were injected bilaterally with 0.5 μL of either lenti-syn-*Bdnf*-eGFP or lenti-syn-eGFP at the rate of 100 nL/min into the VTA (coordinates relative to Bregma: AP −3, ML ±0.5, DV −4.8). Immediately after the surgery, mice were allowed to recover on a heated table (37°C) until becoming fully awake and then returned to their home cages. All mice in the same cage were injected on the same day. Mice were given Carprofen (Rimadyl) chewable tablet (0.5 mg/mouse/day) as analgesics in home cages for 3 days after surgeries.

### Electrophysiology

Mice were anaesthetized via i.p. injection of ketamine/xylazine/acepromazine mixture (69.57 mg/ml; 4.35mg/ml; 0.87mg/ml), then perfused with ice-cold slicing-aCSF consisting of 92 mM N-methyl-d-glucose (NMDG), 2.5 mM KCl, 1.25 mM NaH_2_PO_4_, 10 mM MgSO_4_, 20 mM HEPES, 30 mM NaHCO_3_, 25 mM glucose, 0.5 mM CaCl_2_, 5 mM sodium ascorbate and 3 mM sodium pyruvate, oxygenated with 95% O_2_ and 5% CO_2_. The pH and osmolarity were adjusted to 7.3-7.4 and 315–320 mOsm, respectively. The brain was dissected and embedded with 2% agarose made in slice-aCSF, and 350μm transverse slices containing VTA were cut using vibratome (VF310-0Z, Precisionary Instruments, MA). Slices were first incubated in warm (32°C) slicing-aCSF for 30 mins then transferred to holding-aCSF consisting of 92 mM NaCl, 2.5 mM KCl, 1.25 mM NaH_2_PO_4_, 30 mM NaHCO_3_, 20 mM HEPES, 25 mM glucose, 2 mM MgSO_4_, 2 mM CaCl_2_, 5 mM sodium ascorbate and 3 mM sodium pyruvate, oxygenated with 95% O_2_ and 5% CO_2_. The pH and osmolarity were adjusted to 7.3-7.4 and 310–315 mOsm, respectively. The procedure of acute brain slice preparation was modified from our previous study^52^. Electrophysiological recordings were made in a recording chamber mounted on upright microscope (BX51WI, Olympus Optical Co., Ltd, Tokyo, Japan) with continuous perfusion of warm (29–31°C) recording-aCSF consisting of 124 mM NaCl, 2.5 mM KCl, 1.25 mM NaH_2_PO_4_, 24 mM NaHCO_3_, 5 mM HEPES, 12.5 mM glucose, 2 mM MgCl_2_, 2 mM CaCl_2_, oxygenated with 95% O_2_ and 5% CO_2_. The pH and osmolarity were adjusted to 7.3-7.4 and 310–305 mOsm, respectively. VTA neurons were identified through a highspeed camera (ORCA-Flash4.0LT, Hamamatsu Photonics, Shizuoka, Japan) coupled with water immersion objective lens (LUMPLFLN-40xW, Olympus, Tokyo, Japan). All recordings were made through glass pipettes pulled from borosilicate glass capillary (GC150F-10, Warner Instruments, Hamden, CT) with a resistance around 6-9 MΩ when filled with intra-pipette solution consisting of 120 mM potassium gluconate, 5 mM NaCl, 10 mM HEPES, 1.1 mM EGTA, 15 mM Phosphocreatine, 2 mM ATP and 0.3 mM GTP, with pH 7.2–7.3 and osmolality adjusted to 300 mOsm. Recordings were discarded if the membrane potential (Vm) of recorded cell was over −40 mV or action potentials did not overshoot 0 mV. Recorded cells were continuously clamped at −70 to −75mV under current-clamp mode and received one second current injections from −100 to 400 pA with 10pA steps. Input resistance was calculated using the linear relationship of responses upon current injections between −30 to 0 pA and action potential threshold was measured using the first AP firing upon rheobase current injection. All data were collected using a Multiclamp 700B amplifier (Molecular Devices, San Jose, CA) with a lowpass filter at 2 kHz and digitized at 10k Hz through Axon Digidata 1550B interface (Molecular Devices, CA) coupled with Clampex software (Molecular Devices, CA). Data were further analyzed using MATLAB (MathWorks, MA).

### Statistical analysis and graphing

Data are presented as mean ± standard error of the mean (SEM). Statistical significance was taken as *p≤0.05, **p≤0.01, ***p≤0.001, and ****p≤0.0001, as determined by Student’s *t*-test, Holm-Šídák multiple comparison test, or two-way repeated-measure ANOVA, followed by post-hoc multiple comparisons as appropriate. Significant and close-to-significance main effects of factors, interactions, and pair-wise comparisons are labeled on the graphs. Sample sizes (N) were determined based on empirical estimations. Data sorting and statistical analysis were performed in Microsoft Excel, R studio, and GraphPad Prism 9.0 (Graphpad, La Jolla, CA). Graphs are made with GraphPad Prism 9.0. Cartoons are created with BioRender.com and Microsoft PowerPoint.

## Data availability

All data generated or analyzed during this study are included in this manuscript. Any additional materials are available from the corresponding authors on reasonable request.

## Acknowledgments

We thank Erik D. Herzog and Albert A. Davis for sharing their wheel-running facility and providing critical suggestions on our study and for helping with the primary mesencephalon culture, respectively. We also thank members of the Imai lab, particularly Kyohei Tokizane, Cynthia S. Brace, and Brian V. Lananna, and of the Al-Hasani and McCall labs for technical support and valuable discussions. Support was also provided by the Animal Behavior Core (supported in part by the McDonnell Center for Systems Neuroscience and the Taylor Family Institute) and the Hope Center Viral Vectors Core at Washington University School of Medicine. This work was mainly supported by grants to S.I. from the National Institute on Aging (AG037457, AG047902), and the Tanaka Fund at Washington University School of Medicine. J.G.M. was supported by grants from the National Institute of Neurological Disorders and Stroke (NS11789) and the Brain & Behavior Research Foundation (NARSAD YI-28565). H.C.L. was supported by the LIFENAD Fellowship Fund.

## Contributions

H.C.L. and S.I conceived the project. J.G.M. and S.I. acquired funding and supervised the project. H.C.L., K.E.P., C.M.Y., C.C.K., J.G.M. and S.I. designed experiments, analyzed data, and wrote the manuscript. H.C.L. conducted most experiments, H.C.L. and K.E.P. conducted the *Bdnf* overexpression experiment *in vivo*, C.M.Y. conducted the 1-hour LA/OFT, and C.C.K. conducted the VTA electrophysiology. All authors contributed to the final version of the manuscript.

