## Supplementary Information for "Aging reduces motivation through decreased *Bdnf* expression in the ventral tegmental area"

Campus Box 8103

660 South Euclid Avenue

St. Louis, MO 63110

Tel (Office): 314-362-7228

Fax (Departmental): 314-362-7058

**Jordan G. McCall, Ph.D.**

Associate Professor

Department of Anesthesiology

Center for Clinical Pharmacology

Pain Center

Washington University School of Medicine

Campus Box 8054

660 South Euclid Avenue

St. Louis, MO 63110

Tel (Office): 314-446-8157

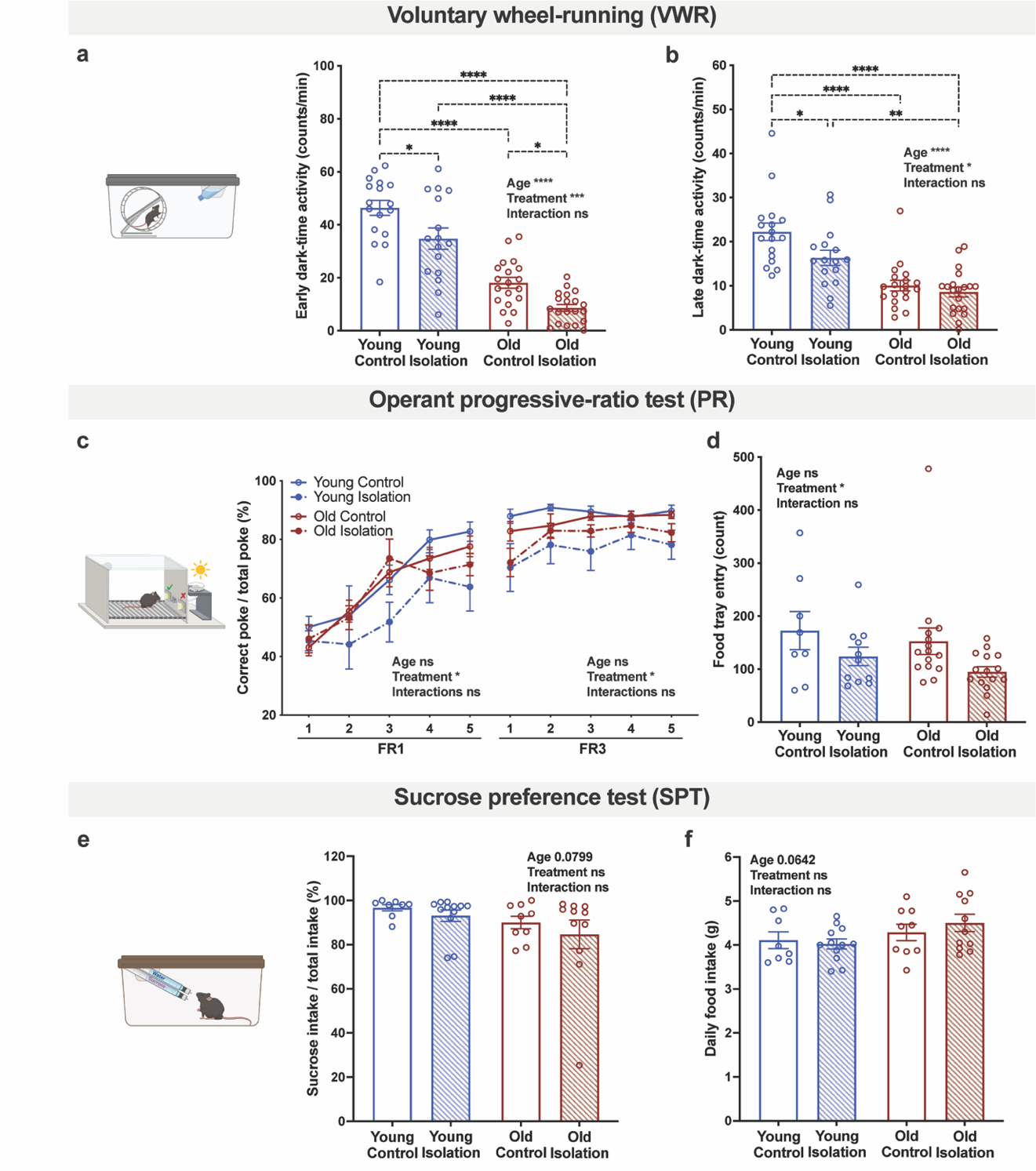

**Extended Data Fig. 1. Chronic social isolation and aging differentially impair motivational behaviors and induce depressive-like behaviors**

Average VWR activity during (**a**) the early dark phase (ZT12-18) and (**b**) the late dark phase (ZT18-0). (**c**) Percentage of correct nose-pokes over total nose-pokes during the FR1 and FR3 training periods. (**d**) The number of entries into the food tray in the PR test. Percentage of sucrose solution intake over total intake (**e**) and daily food intake (**f**) during the test days of SPT. Data were analyzed by 3-way ANOVA (**c**) and 2-way ANOVA (**a**, **b**, **d-f**) with *post-hoc* multiple comparisons. n=7-20. Error bars indicate SEM. Significant pair-wise comparisons and main effects of factors and interactions are labeled on the graphs: *p≤0.05; **p≤0.01; ***p≤0.001; ****p≤0.0001.

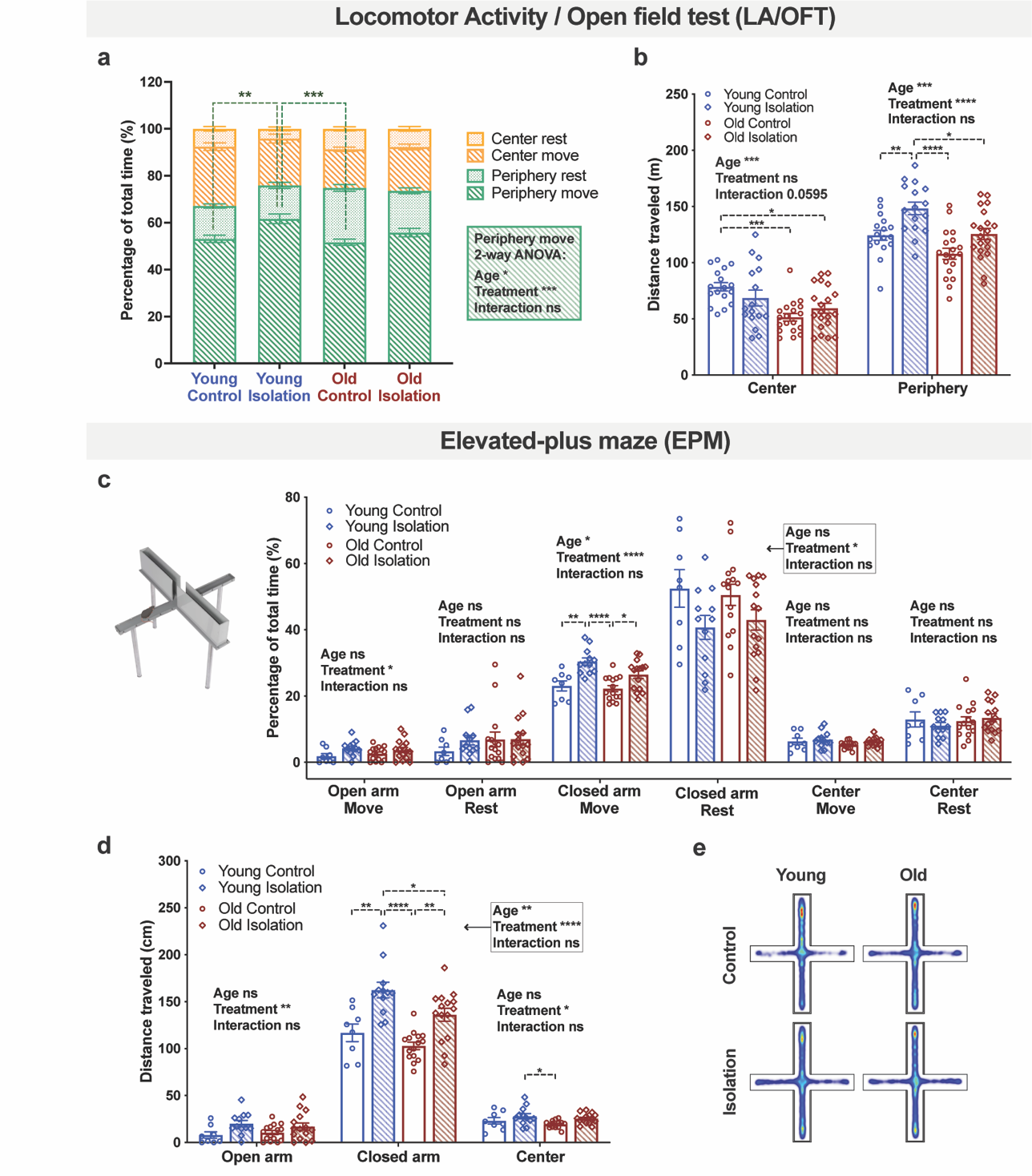

**Extended Data Fig. 2. Chronic social isolation increases locomotor activities in the safe areas of LA/OFT and EPM**

Percentage of time spent moving or resting in the center and peripheral areas (**a**) and total distance traveled in the center and peripheral areas (**b**) in the 1-hour locomotor activity and open field test. Percentage of time spent moving or resting on the open arms, the closed arms, and the center (**c**), and average distance traveled per minute in different areas (**d**) in the elevated elevated-plus maze test. (**e**) Group averaged heat maps of the EPM. Data were analyzed by 2-way ANOVA with *post-hoc* multiple comparisons. n=7-20. Error bars indicate SEM. Significant pair-wise comparisons and main effects of factors and interactions are labeled on the graphs: *p≤0.05; **p≤0.01; ***p≤0.001; ****p≤0.0001.

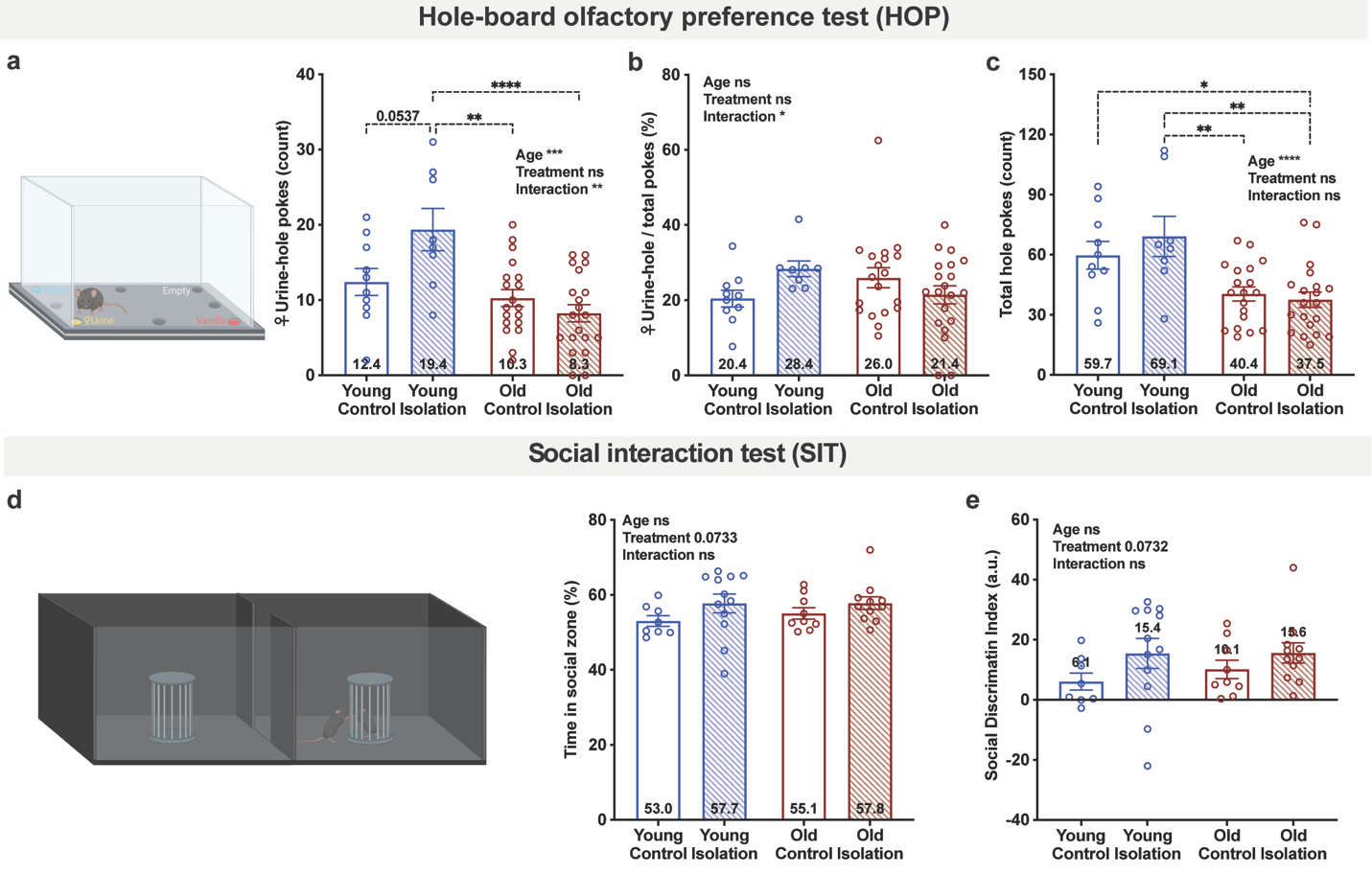

**Extended Data Fig. 3. Chronic social isolation increases preference for social cue only in young mice**

Total number and percentage of pokes into the female urine-containing hole (**a**, **b**) and total number of all pokes (**c**) in the hole-board olfactory preference test. Percentage of time spent in the social zone in the social interaction test (**d**) and calculated social discrimination index. Data were analyzed by 2-way ANOVA with *post-hoc* multiple comparisons. n=8-20. Error bars indicate SEM. Significant pair-wise comparisons and main effects of factors and interactions are labeled on the graphs: *p≤0.05; **p≤0.01; ***p≤0.001; ****p≤0.0001.

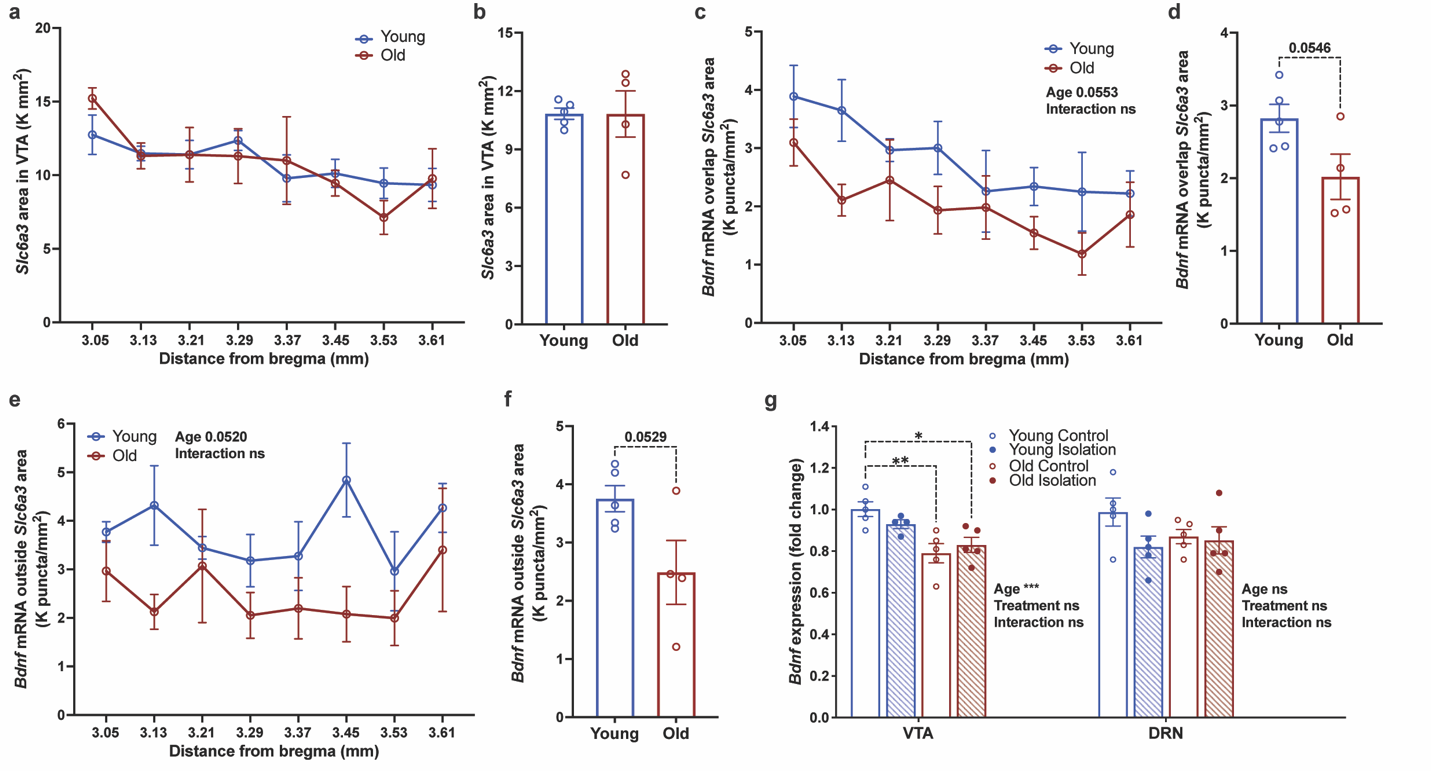

**Extended Data Fig. 4. *Bdnf* gene expression is decreased in the ventral tegmental area (VTA) of aged mice and trending low in the dorsal raphe nucleus (DRN) of young isolated mice**

FISH (RNAscope) analysis of *Bdnf* and Slc6a3 in VTA, quantified as the area of the *Slc6a3* signals (**a**, **b**) and the density of *Bdnf* mRNA signal puncta both overlapping with (**c, d**) and outside of (**e, f**) the *Slc6a3* signals in VTA across 8 bregma levels (80μm interval from -3.05). (**g**) qRT-PCR analysis of *Bdnf* expression in laser-capture-microdissected VTA and DRN samples from young and old, control and isolated mice. Data were analyzed by 2-way ANOVA with *post-hoc* multiple comparisons (**a**, **c**, **e, g**) and student’s *t*-test (**b**, **d, f**). n=4-5. Error bars indicate SEM. Significant pair-wise comparisons, main effects of factors and interactions *p≤0.05; **p≤0.01; ***p≤0.001; ****p≤0.0001.

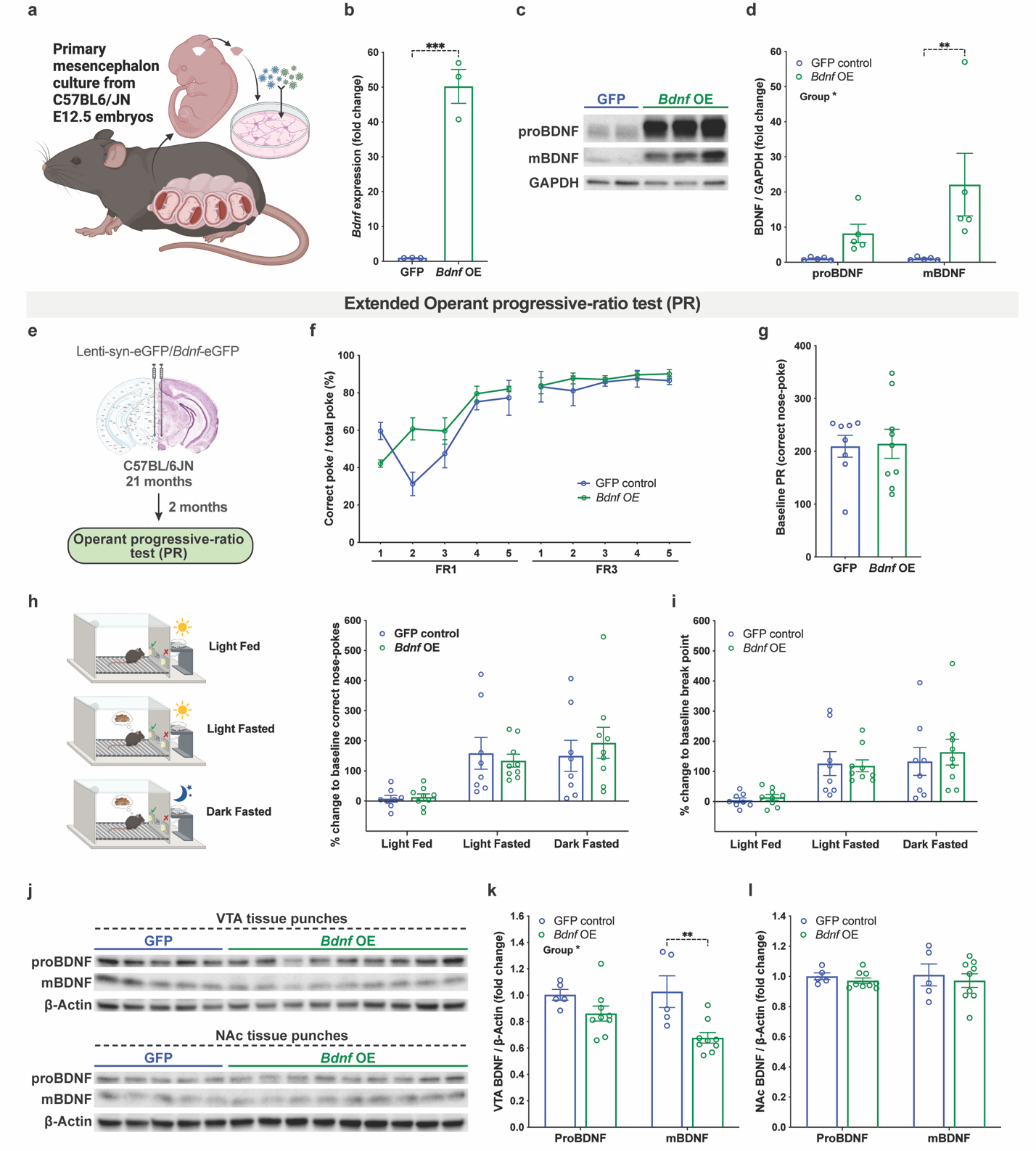

**Extended Data Fig. 5. VTA *Bdnf* overexpression by lentivirus did not increase BDNF protein levels in vivo or alter PR performance in old mice**

(**a**) Schematics illustrating the workflow of virus validation in primary mesencephalon culture. (**b**) qRT-PCR analysis of *Bdnf* mRNA levels in GFP vs *Bdnf*-GFP virus infected primary culture. Representative western blots (**c**) and quantifications (**d**) of proBDNF and mature BDNF (mBDNF) levels normalized to GAPDH signals in primary culture. (**e**) A scheme illustrating the experimental design. (**f**) Percentage of correct nosepokes over total nosepokes during the FR1 and FR3 training period. (**g**) Correct nosepokes in baseline PR (mean of 2 tests) in the extended PR. Correct nosepokes (**h**) and break points reached (**i**) normalized to individual baseline PR under different experimental conditions in the extended PR. Western blots (**j**) and quantifications (**k, l**) of proBDNF and mBDNF levels normalized to β-ACTIN signals in punched VTA and NAc of GFP vs. *Bdnf*-GFP virus injected mice after behaviors tests. Data were analyzed by 2-way ANOVA with *post-hoc* multiple comparisons (**d**, **f**, **h**, **i**, **k, l**), student’s *t*-test (**b**, **g**). n=3-9. Error bars indicate SEM. Significant pair-wise comparisons and main effects of factors and interactions are labeled on the graphs: *p≤0.05; **p≤0.01; ***p≤0.001; ****p≤0.0001.

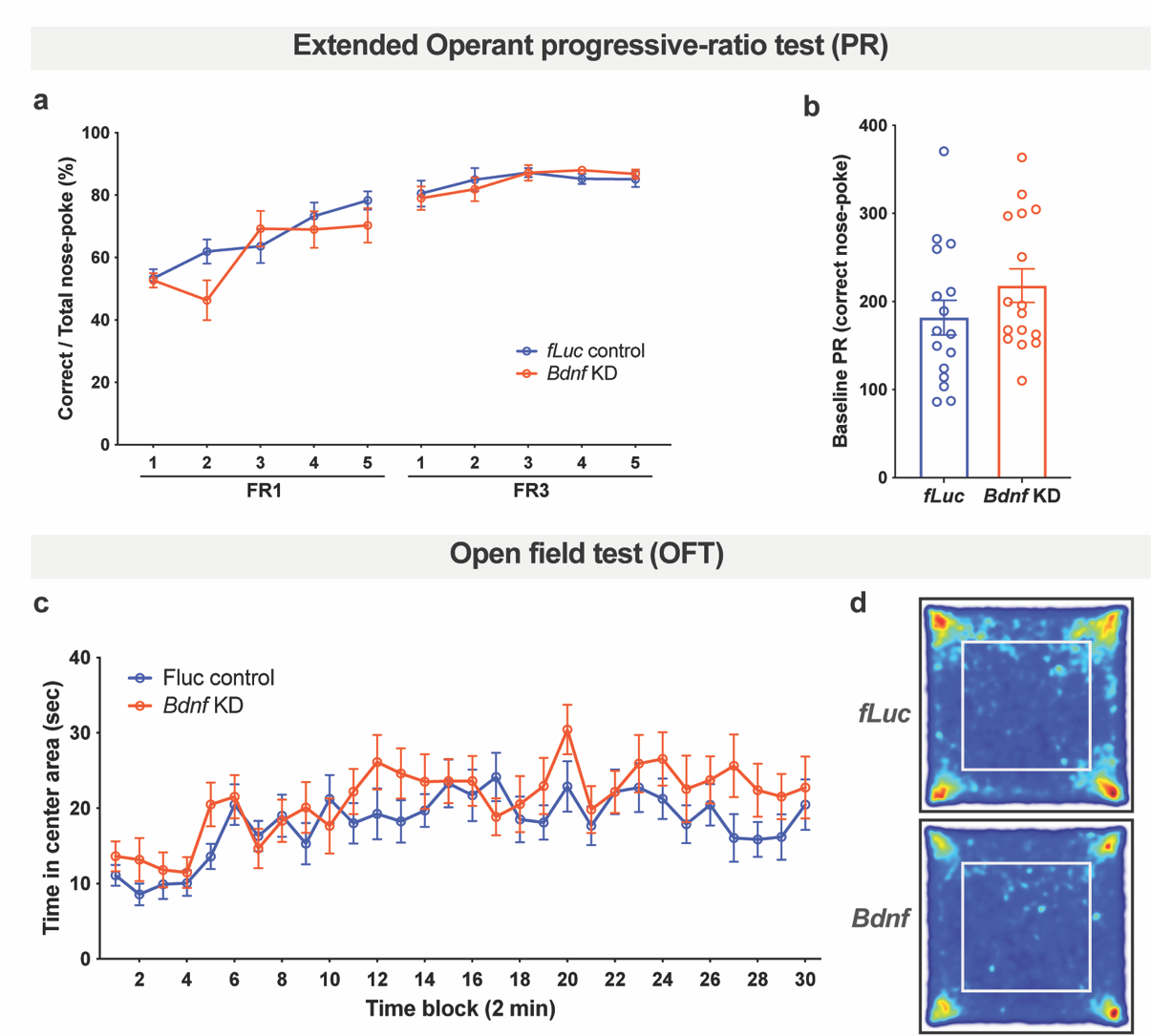

**Extended Data Fig. 6. VTA *Bdnf* KD in young mice partially recapitulates the age-associated reduction in palatable food motivation, not affecting other age-related phenotypes**

(**a**) Percentage of correct nose-pokes over total nose-pokes during the FR1 and FR3 training period. (**b**) Correct nose-pokes in baseline PR (mean of 2 tests) in the extended PR. (**c**) Percentage of time spent in the center area across 2-min time blocks in the open field test. (**d**) Group averaged heat maps of the OFT. Data were analyzed by 2-way ANOVA with *post-hoc* multiple comparisons (**a**, **c**) and student’s *t*-test (**b**). n=8-18. Error bars indicate SEM. Significant pair-wise comparisons, main effects of factors and interactions *p≤0.05; **p≤0.01; ***p≤0.001; ****p≤0.0001.

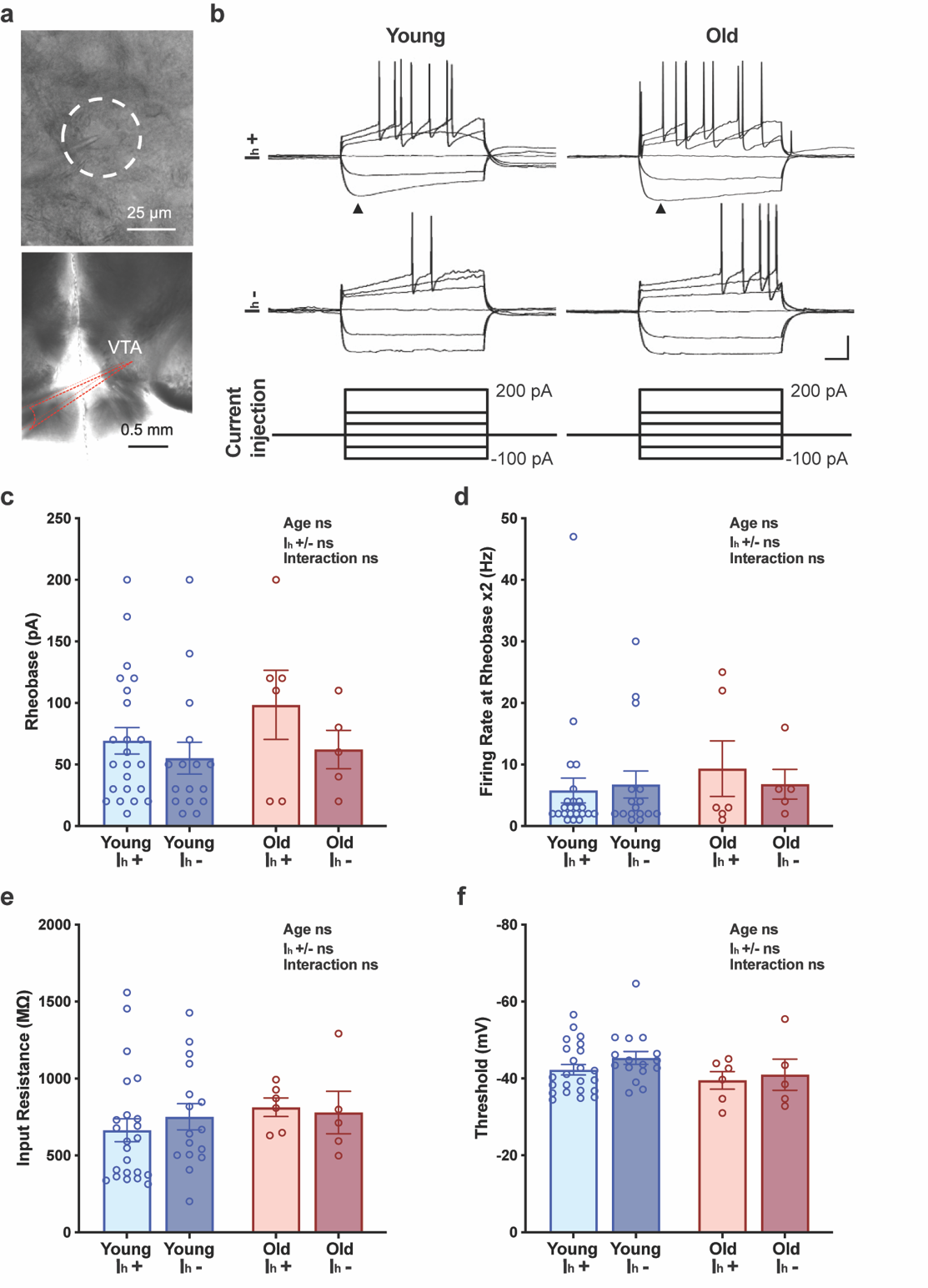

**Extended Data Fig. 7. Neuronal excitability is unchanged in aged VTA neurons**

(**a**) Live DIC images in high (40x) and low (5x) magnifications demonstrating an electrophysiological recording on VTA neurons. (**b**) Representative current clamp recording traces showing the responses to current injections from -100 to 200 pA in young (3-4 months) and old (19-20 months) mice. The presence of hyperpolarization-activated cation current (I_h_) (marked by triangles) upon hyperpolarized current injection was used to identify putative dopaminergic (I_h_ +) and non-dopaminergic (I_h_ -) neurons. Assessments of neuronal excitability do not show significant differences between neurons recorded from young and old mice with measurements including rheobase (**c**), firing rate upon 2 times of rheobase (**d**), input resistance (**e**), and threshold (**f**). Data were analyzed by 2-way ANOVA with *post-hoc* multiple comparisons. n=5-23. Error bars indicate SEM.
